# CT Radiomic Signatures of Neutrophil Extracellular Traps in Ischemic Stroke Thrombi

**DOI:** 10.1101/2025.08.11.669792

**Authors:** Briana A. Santo, Tatsat R. Patel, Seyyed Mostafa Mousavi Janbeh Sarayi, Kerry E. Poppenberg, Sarah Balghonaim, Alexandria Scotti, TaJania D. Jenkins, Elad I. Levy, Adnan H. Siddiqui, John Kolega, Vincent M. Tutino

**Affiliations:** Canon Stroke and Vascular Research Center; Buffalo, NY, USA; Department of Pathology and Anatomical Sciences, Buffalo, NY, USA; Department of Neurosurgery, Buffalo, NY, USA; Department of Mechanical and Aerospace Engineering, University at Buffalo, Buffalo, NY, USA

**Keywords:** Ischemic Stroke, Thrombosis, Mechanics, Immunology, Blood, Thrombectomy

## Abstract

**Background:** Radiomic and transcriptomic analyses have independently identified features linked to mechanical thrombectomy (MT) outcomes in acute ischemic stroke (AIS). In this study, we integrate paired radiomic and transcriptomic profiling of AIS clots to identify Neutrophil Extracellular Trap (NET) enrichment as a predictor of first-pass MT success. We further assess the potential to non-invasively detect NET enrichment using pre-thrombectomy CT imaging.

**Methods:** We performed radiomic and transcriptomic analysis of 32 stroke clots retrieved by MT (n=16 each of modified first pass [mFP] success and failure). Clots were segmented from pre-MT CTA and nCCT scans and radiomic features (RFs) were extracted using pyRadiomics. Normality, equal variance, and two-sample testing were completed to identify which RFs were significantly different between mFP outcomes. Differentially expressed genes (DEGs) were identified between transcriptomes of mFP success and failure using the criteria of logFC≥1.5 and q<0.05. A NET enrichment score was computed from expression data and correlated with RFs to identify a RF signature predictive of NET enrichment. Immunofluorescence (IF) staining was completed on retrieved clot tissue to provide ground truth labeling of NETs.

**Results:** 44 DEGs were identified between mFP outcomes. From ontology analysis, NET Formation, Neutrophil Degranulation, and the NET Signaling Pathway were among the most enriched terms in the mFP failure group, with related genes downregulated in the mFP success group. 40 RFs were significantly different between mFP outcomes. Of these, 6 were found to be correlated with and predictive of clot NET enrichment. IF quantification validated that transcriptomic NET signatures accurately reflected NET presence within clot tissues.

**Conclusion:** Our findings indicate that NET enrichment within thrombi is associated with reduced mFP success, and that radiomic features extracted from pre-thrombectomy CT imaging can serve as non-invasive biomarkers of clot NET content. This radiomic signature may aid in pre-procedural decision-making, including thrombolytic therapy planning and thrombectomy device selection.

## Introduction

Acute ischemic stroke (AIS) poses a significant global health burden, necessitating rapid and effective interventions to mitigate significant morbidity and mortality.^1^ Mechanical thrombectomy (MT) has emerged as a pivotal therapeutic approach, offering a lifeline for patients afflicted by large vessel occlusions.^2^ The success of MT, particularly on the first pass (FP), represents a critical milestone in achieving optimal outcomes.^3^ To best predict likelihood of MT success, studies have focused on learning what properties of a clot make it challenging to remove mechanically. A large body of work has been dedicated to investigating whether clot radiomics from pre-thrombectomy CT or clot composition (percentage red blood cell [RBC], white blood cell [WBC], and fibrin platelet [FP] meshwork) determined from digital histology can predict MT FP outcome.^4–7^ While some findings have had consensus, such as stiffer clots being less amenable to retrieval,^8,9^ reports on composition or which biological components of the clot are important, are inconsistent.^10,11^ It has been suggested that percent composition, alone, is insufficient to decipher the biology of AIS clots because composition overlooks important factors such as the density of FP meshwork, the localization of WBCs along the RBC-FP interface, and the enrichment of pro-thrombotic phenomenon such as Neutrophil Extracellular Traps (NETs).^12,13^

NETs are networks of extracellular DNA, antimicrobial enzymes, and histone that serve a vital antimicrobial function in the body.^14^ However, ischemic conditions such as those in stroke can facilitate rapid NET formation wherein NETs create a hypercoagulable state and reinforce thrombus microstructure. In the *in vitro* clot setting, NETs have been observed to bind fibrin, increase crosslinking, and produce denser microstructure.^15^ The compaction that results increases clot stiffness and resistance to both mechanical disruption (MT) and dissolution (tPA resistance).^16^ Additionally, NET DNA strands with a dense negative charge serve as a scaffold for other blood components (FP, RBC) while enzymes elicit further immune response. In vitro studies have also shown that the addition of DNase I to tPA can help resolve clots into the blood flow, supporting the concept that the NET DNA backbone is a key component altering clot microstructure.

*In vitro* studies have demonstrated that NETs significantly alter the biophysical properties of thrombi.^17,18^ In addition, ongoing clinical trials are investigating whether DNA-targeting therapeutics can enhance early reperfusion and improve outcomes in patients with large vessel occlusion stroke. These agents have demonstrated efficacy in degrading NETs and improving outcomes in conditions such as acute respiratory distress syndrome and hypercoagulable states.^19^ Given this, quantifying NET enrichment in AIS thrombi may help identify patients who would benefit most from adjunctive therapies (e.g., tPA combined with DNase) or guide the selection of mechanical thrombectomy (MT) devices. Currently, NET detection in stroke clots relies solely on post-retrieval immunofluorescence staining, and no validated imaging-based biomarkers exist for pre-procedural assessment. To address this gap, we analyzed NET enrichment in AIS thrombi and investigated its association with radiomic features derived from pre-thrombectomy CT imaging. If validated in larger cohorts, a radiomic signature of NET content could enable personalized treatment strategies within the critical stroke intervention window.

## Methods

### Patient Data

This study was approved by our institute’s institutional review board (study 00002092). All methods were performed in accordance with the approved protocol, and written informed consent was obtained for all subjects. Clot samples and imaging were collected as previously described from patients receiving MT by either stent retriever, aspiration, or a combined therapy between November 2018 and November 2020 at our institution.^13^ Patients considered for this study met the following criteria: 1) They had undergone MT for acute ischemic stroke, 2) had pretreatment CT available with sufficient image quality and an identifiable clot, 3) had a retrieved clot of sufficient size and quality for microCT and histology, and 4) did not receive intra-arterial tPA. MT first pass outcomes were defined using the modified first pass effect (mFPE) definition of thrombectomy success, which in the context of endovascular thrombectomy is defined as complete or near-complete reperfusion (Thrombolysis in Cerebral Infarction [TICI] score of 2b/2c/3) on the first attempt with a thrombectomy device.^3^

### Clot mRNA Extraction and Differential Expression Analysis

Clot transcriptomes were obtained from those analyzed in our previous work that established a pipeline for obtaining useful RNA from AIS blood clots.^20^ In brief, clot samples retrieved by MT and stored in RNALater were subjected to RNA extraction using the Chemagen magnetic bead extraction protocol on a PerkinElmer Chemagic 360. A total of 48 samples were used for RNA sequencing based on sequencing quality metrics. In this study, a subset of n=32 clot transcriptomes for which histology was also available was examined to determine if expression profiles were unique to MT first pass outcomes. Differential gene expression analysis was completed between patient samples that did or did not achieve mFPE using a Fischer’s Exact Test in edgeR, as previously described.^21,22^ Significant differentially expressed genes (DEGs) were identified using the following criteria: 50% minimum expression among samples, absolute logFC≥1.5, and q<0.05 after false discovery correction by the Benjamini-Hochberg (BH) procedure. All gene candidates were plotted as a volcano plot, and DEGs were denoted as colored dots based on whether they were up-or down-regulated. Using *pheatmap* in R, DEGs were visualized as a heatmap and hierarchically clustered based on Euclidean distance between observations.^23^

### Ontology Analyses and Calculation of Clot NET Expression

To understand the biology of mFPE-associated DEGs, we performed gene set enrichment analyses using *ClusterProfiler*^*24*^ and QIAGEN Ingenuity Pathway Analysis (IPA, QIAGEN Inc., https://digitalinsights.qiagen.com/). For *ClusterProfiler*, we queried the KEGG^25^ and ReactomePA databases using our DEG set.^26^ For IPA, we specifically analyzed enrichment of the disease and biological function terms (terms that had q-value<0.05 and 3 or more input genes were considered).

Based on the results of ontology analysis, we further sought to compute the per sample enrichment of NET formation based on clot transcriptomes. To achieve this, we pulled a comprehensive list of all genes defining the KEGG “Neutrophil Extracellular Trap Formation” term (hsa04613) using *keggGet*.^27^ We then computed the ssGSEA (Single Sample Gene Set Enrichment Analysis) score for each sample using *gsva*^28^ (Gene Set Variation Analysis) and normalized the set of scores between zero and one as recommended. High and low NET enrichment groups were stratified by the median NET score. All analysis was completed in R v4.4.2.

### Immunofluorescence Labeling and Imaging of Neutrophil Extracellular Traps in Retrieved Clots

To provide ground truth values of clot NET enrichment, we completed immunofluorescence staining, whole-slide imaging, and whole-slide quantification of MT-retrieved clot tissues. Clot histological processing and immunofluorescence staining were completed as previously described.^29^ In brief, formalin-fixed paraffin embedded tissues were sectioned at 4μm thick and mounted on clear glass slides. Indirect immunofluorescence labeling of citrullinated histone was completed using Rabbit Anti Citrated Histone (ab281584, 1:200μg/mL, Abcam, Cambridge, UK), donkey blocking serum, and Donkey Anti Rabbit AF647 secondary antibody (ab150075, 1:200μg/mL, Abcam, Cambridge, UK). Slides were then DAPI counterstained, and #1.5 coverslips were applied. Whole-slide fluorescence images were acquired at 20X on a Leica DM6B Fluorescence Microscope (Leica Biosystems, Lincolnshire, IL). The DAPI and Y5 filter cubes were used to image DAPI and AF647 (citrullinated histone), respectively (exposure time was 100-200ms for DAPI and 400ms for AF647).

The percentage of citrullinated histone, an established measure of clot NET enrichment,^30–32^ was then computed as follows. The blue (DNA) and red (citrullinated histone) channels of fluorescence WSIs were extracted, and a top-hat filter was applied to each to correct for uneven illumination. Global mean-based thresholds were applied to each channel to segment true-positive signal from background and produce binary masks for DNA and citrullinated histone. The intersection of DNA and citrullinated histone (DNAxCitHis) regions was found by taking the intersection (logical AND) of the respective masks. This region represented the NET-positive region. The area of citrullinated histone within the NET-positive area was indexed to the total clot area to yield %Citrullinated Histone.

### CT Image Analysis

For all patients, CT was performed on an Aquilion ONE scanner (Canon Medical Systems). In this study, CTA and nCCT images with the lowest thickness (0.5 mm) and highest resolution were used for analysis. As previously described, 3D Slicer was used to register CTA and nCCT images, manually segment and reconstruct patient vasculatures, and isolate the clot region of interest^33^. Clot annotation was completed by two experienced annotations, and inter-rater agreement was assessed by Dice Coefficient. PyRadiomics, an open-source Python package, was used to extract clot morphology and textural features from segmented clot regions in both the CTA and nCCT images (bin size=25, no resampling). In addition, textural feature ratios were computed by dividing the value of a given radiomic from CTA by its value in nCCT, yielding radiomic feature ratios (CTA:nCCT). Textural features included first order statistics (FOS), gray-level run length matrix (GLRLM), gray-level dependence matrix (GLDM), gray-level co-occurrence matrix (GLCM), and gray-level size zone matrix (GLSZM). Formulae for RFs are available in the PyRadiomics documentation (https://pyradiomics.readthedocs.io/). Combined, CTA, nCCT, and CTA:nCCT clot radiomic features totaled 293 features (93 per modality, 93 ratios, and 14 shape). Feature values were normalized between zero and one.

### Statistical Analysis

Statistical analysis was completed to identify radiomic features significantly different between first pass outcomes, as well as to assess correlation between radiomics and DEGs and between radiomics and NET scores. Normality was assessed using the Anderson Darling Test Statistic. Correlation analysis was completed using Pearson (parametric) or Spearman (nonparametric) correlation. Given our sample size, radiomics associated with clot NET score had to meet two criteria: p<0.1 and R>0.3. To assess the predictive power of radiomics for clot NET scores, a multivariable logistic regression model was fit using correlated radiomics as independent variables and NET score as the dependent variable. For the model, a ROC curve was produced and the area under the curve (AUC) was reported. All statistical analysis was completed in R v4.4.2 and statistical significance was considered at α<0.05.

## Results

### Summary of Patient Population

A summary of characteristics for the 32 patients (59.3% female) included in this study is provided in Table 1. The mean (± standard deviation) age of patients was 71.3±14.7 years, and NIHSS at presentation was 16.2±7.2. In our cohort, 3.1% of strokes were treated with stent retriever thrombectomy, 65.6% with aspiration, and 31.2% with combination therapy (Solumbra). Occlusions were primarily located on the right (65.6%); 9.4% of clots were located in the ICA, 25% at the ICA/M1 junction, 37.5% in the MCA M1 region, 18.7% in the MCA M2 region, and 9.4% at the basilar. In all, 50% of thrombectomy procedures achieved mFPE.

**Table 1:**
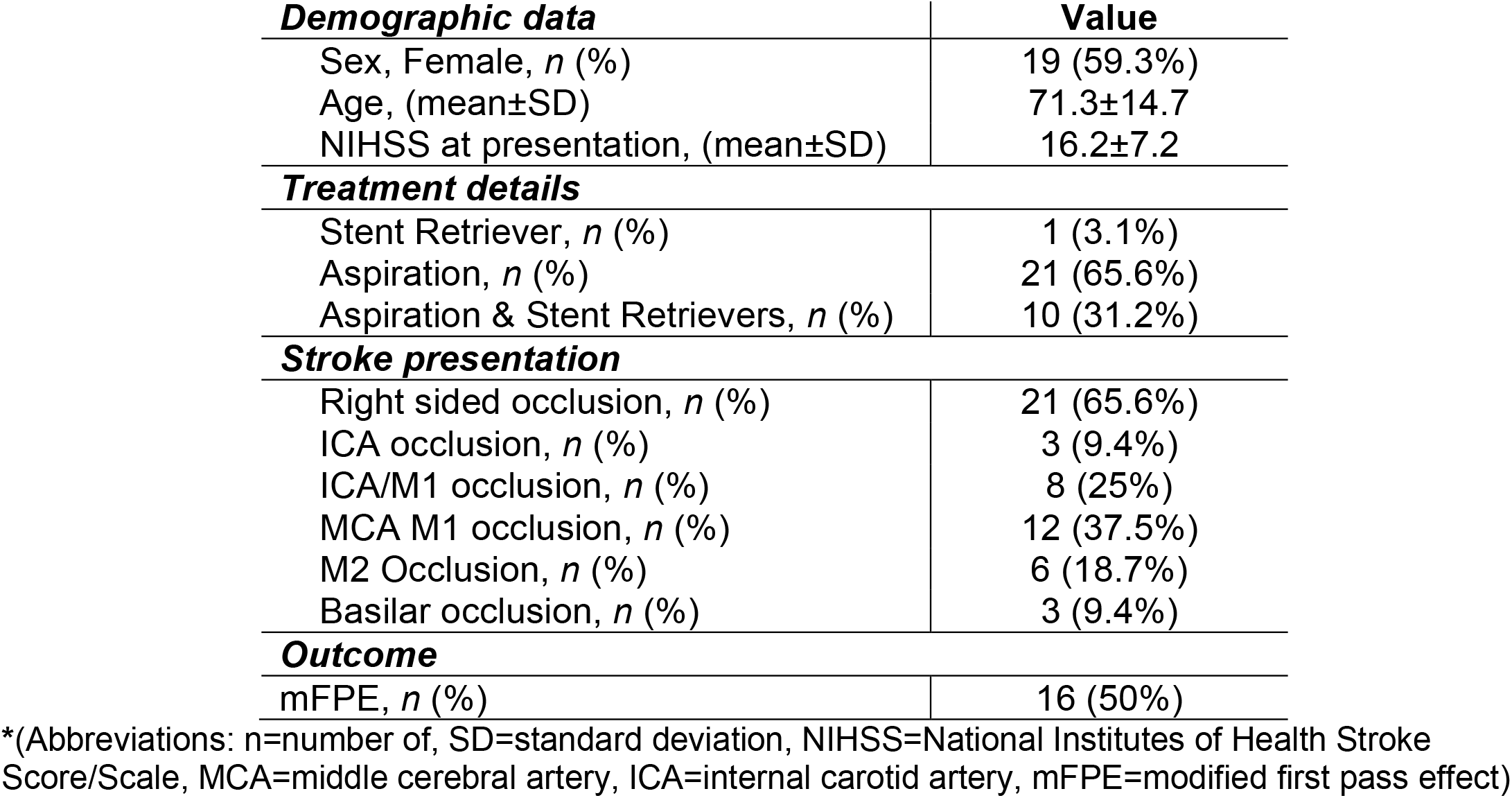
Baseline patient characteristics.*

| ***Demographic data*** | **Value** |
| --- | --- |
| Sex, Female, *n* (%) | 19 (59.3%) |
| Age, (mean±SD) | 71.3±14.7 |
| NIHSS at presentation, (mean±SD) | 16.2±7.2 |
| ***Treatment details*** |  |
| Stent Retriever, *n* (%) | 1 (3.1%) |
| Aspiration, *n* (%) | 21 (65.6%) |
| Aspiration & Stent Retrievers, *n* (%) | 10 (31.2%) |
| ***Stroke presentation*** |  |
| Right sided occlusion, *n* (%) | 21 (65.6%) |
| ICA occlusion, *n* (%) | 3 (9.4%) |
| ICA/M1 occlusion, *n* (%) | 8 (25%) |
| MCA M1 occlusion, *n* (%) | 12 (37.5%) |
| M2 Occlusion, *n* (%) | 6 (18.7%) |
| Basilar occlusion, *n* (%) | 3 (9.4%) |
| ***Outcome*** |  |
| mFPE, *n* (%) | 16 (50%) |

\*(Abbreviations: n=number of, SD=standard deviation, NIHSS=National Institutes of Health Stroke Score/Scale, MCA=middle cerebral artery, ICA=internal carotid artery, mFPE=modified first pass effect)

### Transcriptomic Signatures Associated with Modified First Pass Effect

To identify clot transcriptomic signatures associated with mFPE, we performed differential gene expression analysis comparing patients who achieved mFPE to those who did not. A total of 44 differentially expressed genes (DEGs) were identified, with 1 gene upregulated and 43 downregulated in the mFPE success group (Figure 1A–B, Table 2). The sole gene with significantly increased expression in successful first pass cases was *CRIP2* (q = 0.031). The top three DEGs with significantly decreased expression were *MMP8* (q < 0.001), *MS4A3* (q < 0.001), and *TCN1* (q = 0.004).

**Table 2:**
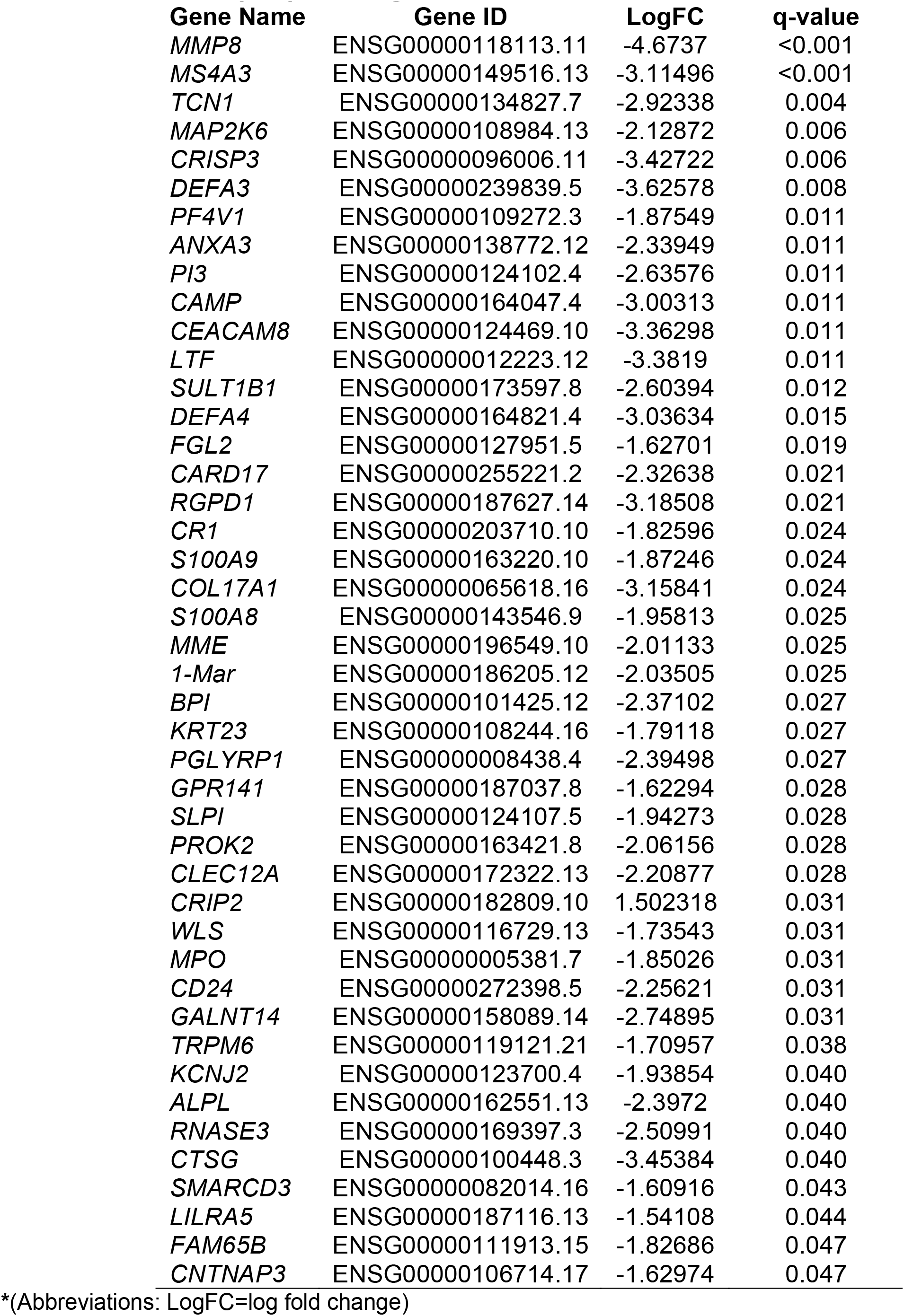
Differentially expressed genes between mFPE outcomes.*

**Figure 1:**
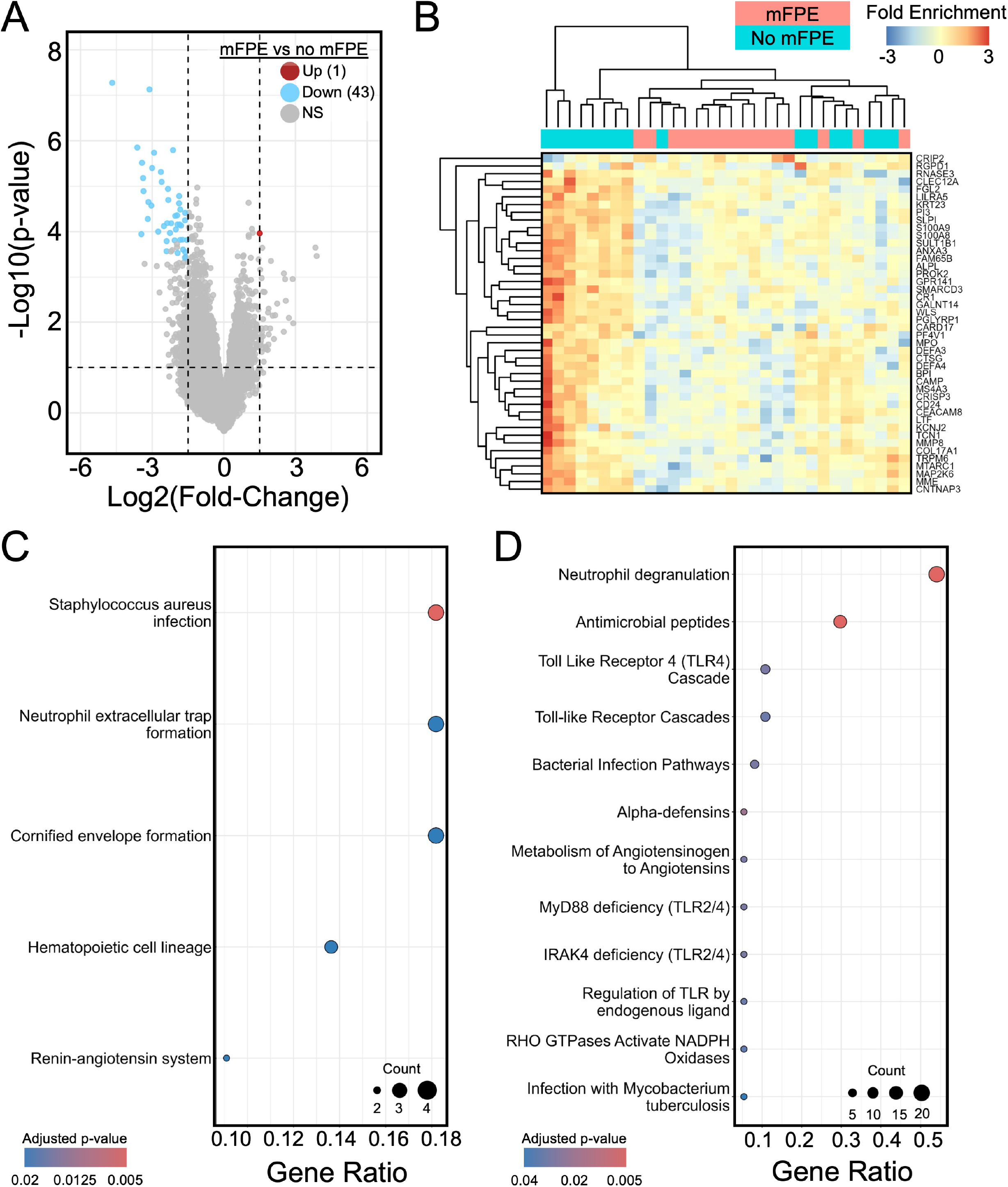
Clot gene expression differences between mFPE success and failure thrombectomy attempts. **A)**. A volcano plot of differentially expressed genes shows 43 genes were downregulated, and one gene was upregulated in mFPE successes. **B)**. Hierarchical clustering of differentially expressed genes shows separation between mFPE successes and failures. **C)**. KEGG analysis of 44 differentially expressed genes identified pathways enriched in mFPE failures, including neutrophil extracellular trap formation **D)**. ReactomePA analysis of fourty-four differentially expressed genes identified pathways enriched in mFPE failures including neutrophil degranulation. (Abbreviations: mFPE=modified first pass effect; NS=not significant)

### Neutrophil Extracellular Trap Formation and Signaling are Associated with Thrombectomy Failure

To better understand the biological significance of DEGs associated with mFPE, we completed gene set enrichment analysis (GSEA) with KEGG, ReactomePA, and IPA databases. From KEGG, the two most enriched pathways were *S. aureus* infection (q=0.005) and Neutrophil extracellular trap formation (q=0.018) (Figure 1C). ReactomePA also emphasized neutrophil activity, with top terms of Neutrophil degranulation (q<0.001) and Antimicrobial peptides (q<0.001) (Figure 1D). When the ssGSEA score for KEGG Neutrophil extracellular trap formation was plotted for each of the mFP successes and failures, increased NET formation in mFP failures was observed (Figure S1A). GSEA with IPA further supported these findings, with the Neutrophil Extracellular Trap Signaling Pathway identified as the most enriched pathway in mFP failures (negative value in mFP successes) (Figure S1B).

### Clot Radiomics are Significant Predictors of mFPE

We next sought to identify CT radiomic signatures associated with mFP outcomes. Inter-rater assessment of clot annotations by two experienced annotators demonstrated high agreement, with a mean dice coefficient of 0.80±0.09 (Figure 2A). A total of 40 RFs computed from CTA and nCCT clot segmentations were significantly different between mFP outcomes (Table 3). Of these RFs, 34 were computed from CTA, 1 from nCCT, and 5 from the ratio of CTA to nCCT (Figure 2B). Overall, three RFs were common to the CTA and CTA:nCCT ratio analyses (Figure 2B). These RFs were all derived from the gray-level co-occurrence matrix (GLCM) and included Autocorrelation, Joint energy, and Maximum probability. Two significant RFs derived from CTA, FOS 90^th^ Percentile and GLCM Joint Energy, were visualized as heatmaps and superimposed on pre-thrombectomy CTA for further interpretation. Here, a significant increase in FOS 90^th^ Percentile (p=0.021) was observed in mFP successes, suggesting that clots with a higher 90^th^ Percentile intensity value are more amenable to retrieval (Figure 2C). In comparison, a significant increase in GLCM Joint energy (p=0.017) was observed in mFP failures, suggesting that clots with greater variation in intensity values, and thus textural complexity, are more likely to result in mFP failure (Figure 2D).

**Table 3:** Radiomics significantly different between mFPE outcomes.*

| **Radiomic Feature** | **p-value** |
| --- | --- |
| FOS 90<sup>th</sup> Percentile, CTA | 0.022 |
| FOS Energy, CTA | 0.033 |
| FOS InterquartileRange, CTA | 0.018 |
| FOS Kurtosis, Ratio | 0.019 |
| FOS MeanAbsoluteDeviation, CTA | 0.034 |
| FOS Mean, CTA | 0.047 |
| FOS RobustMeanAbsoluteDeviation, CTA | 0.022 |
| FOS RootMeanSquared, CTA | 0.038 |
| FOS Skewness, Ratio | 0.004 |
| FOS TotalEnergy, CTA | 0.023 |
| FOS Uniformity, CTA | 0.048 |
| FOS Variance, CTA | 0.036 |
| GLCM Autocorrelation, CTA | 0.017 |
| GLCM Autocorrelation, Ratio | 0.031 |
| GLCM ClusterProminence, CTA | 0.029 |
| GLCM ClusterTendency, CTA | 0.028 |
| GLCM Contrast, CTA | 0.049 |
| GLCM DifferenceAverage, CTA | 0.046 |
| GLCM DifferenceEntropy, CTA | 0.033 |
| GLCM Id, CTA | 0.049 |
| GLCM Idm, CTA | 0.047 |
| GLCM Idmn, CTA | 0.033 |
| GLCM InverseVariance, CTA | 0.046 |
| GLCM JointAverage, CTA | 0.019 |
| GLCM JointEnergy, CTA | 0.017 |
| GLCM JointEnergy, Ratio | 0.037 |
| GLCM JointEntropy, CTA | 0.012 |
| GLCM MaximumProbability, CTA | 0.046 |
| GLCM MaximumProbability, Ratio | 0.042 |
| GLCM SumAverage, CTA | 0.019 |
| GLCM SumEntropy, CTA | 0.025 |
| GLCM SumSquares, CTA | 0.028 |
| GLDM GrayLevelVariance, CTA | 0.038 |
| GLDM HighGrayLevelEmphasis, CTA | 0.022 |
| GLRLM GrayLevelVariance, CTA | 0.045 |
| GLRLM HighGrayLevelRunEmphasis, CTA | 0.023 |
| GLRLM LongRunHighGrayLevelEmphasis, CTA | 0.029 |
| GLRLM ShortRunHighGrayLevelEmphasis, CTA | 0.026 |
| Shape Maximum2DDiameterRow, CTA | 0.030 |
| Shape Maximum2DDiameterRow, nCCT | 0.030 |

\*(Abbreviations: FOS=first order statistic; GLCM=gray-level co-occurrence matrix; GLDM=gray-level dependence matrix; GLRLM=gray-level run length matrix)

**Figure 2:**
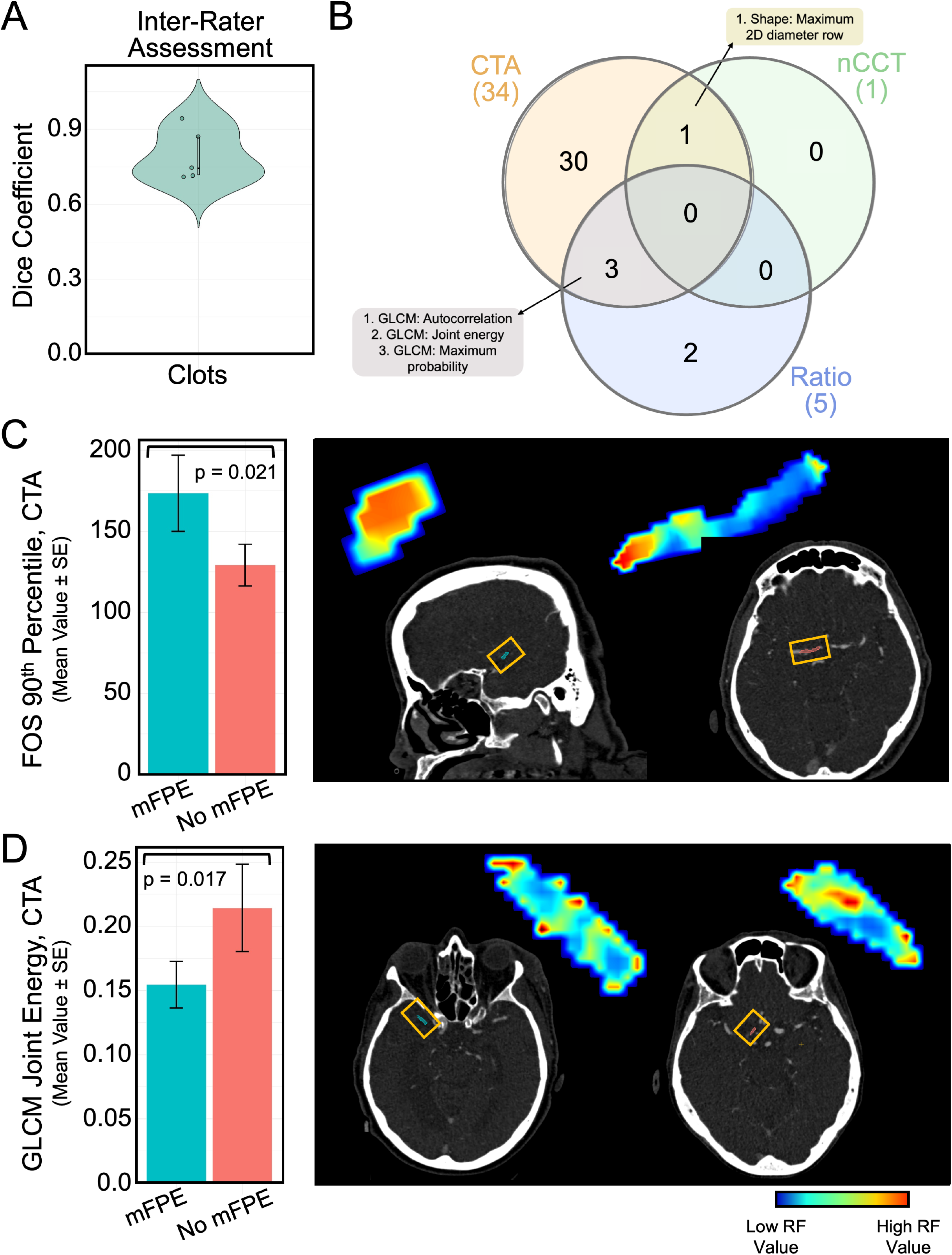
Clot radiomic differences between mFPE success and failure thrombectomy attempts. **A)**. Inter-rater assessment of clot annotations shows high agreement. **B)**. Venn diagram shows significant radiomics common to CTA, nCCT, and ratio of CTA:nCCT feature sets. **C)**. Bar plot shows clot first order statistic 90^th^ percentile measured from CTA as a radiomic significantly different between mFPE success and failure. Radiomic heatmaps visualize the first order statistic 90^th^ percentile in example cases of mFPE success and failure. **D)**. Bar plot shows clot gray-level co-occurrence matrix joint energy measured from CTA as a radiomic significantly different between mFPE success and failure. Radiomic heatmaps visualize the gray-level co-occurrence joint energy in example cases of mFPE success and failure. (Abbreviations: mFPE=modified first pass effect; FOS=first order statistic; GLCM=gray-level co-occurrence matrix)

### Pre-thrombectomy CT Radiomics May Predict Clot Neutrophil Extracellular Trap Enrichment

Having identified both clot transcriptomic and radiomic signatures of mFP outcome, as well as the significance of NET gene set enrichment, we sought to assess whether an association between pre-thrombectomy clot radiomics and NET enrichment could be identified.

#### Validation of Clot NET Enrichment by Immunofluorescence Quantification

After binarizing clot NET scores using the median, a distribution-independent measure, we arrived at 14 clots in both the low and high NET enrichment groups (p<0.001) (Figure 3A). We validated this finding through quantification of ground truth immunofluorescence labeling of NET formation in clots retrieved by MT (Figure 3B). Here, NET formation was quantified as the percentage of citrullinated histone. When the percentage of citrullinated histone was compared between transcriptome-defined low and high NET enrichment clots, we identified a significant difference (p=0.007) (Figure 3C).

**Figure 3:**
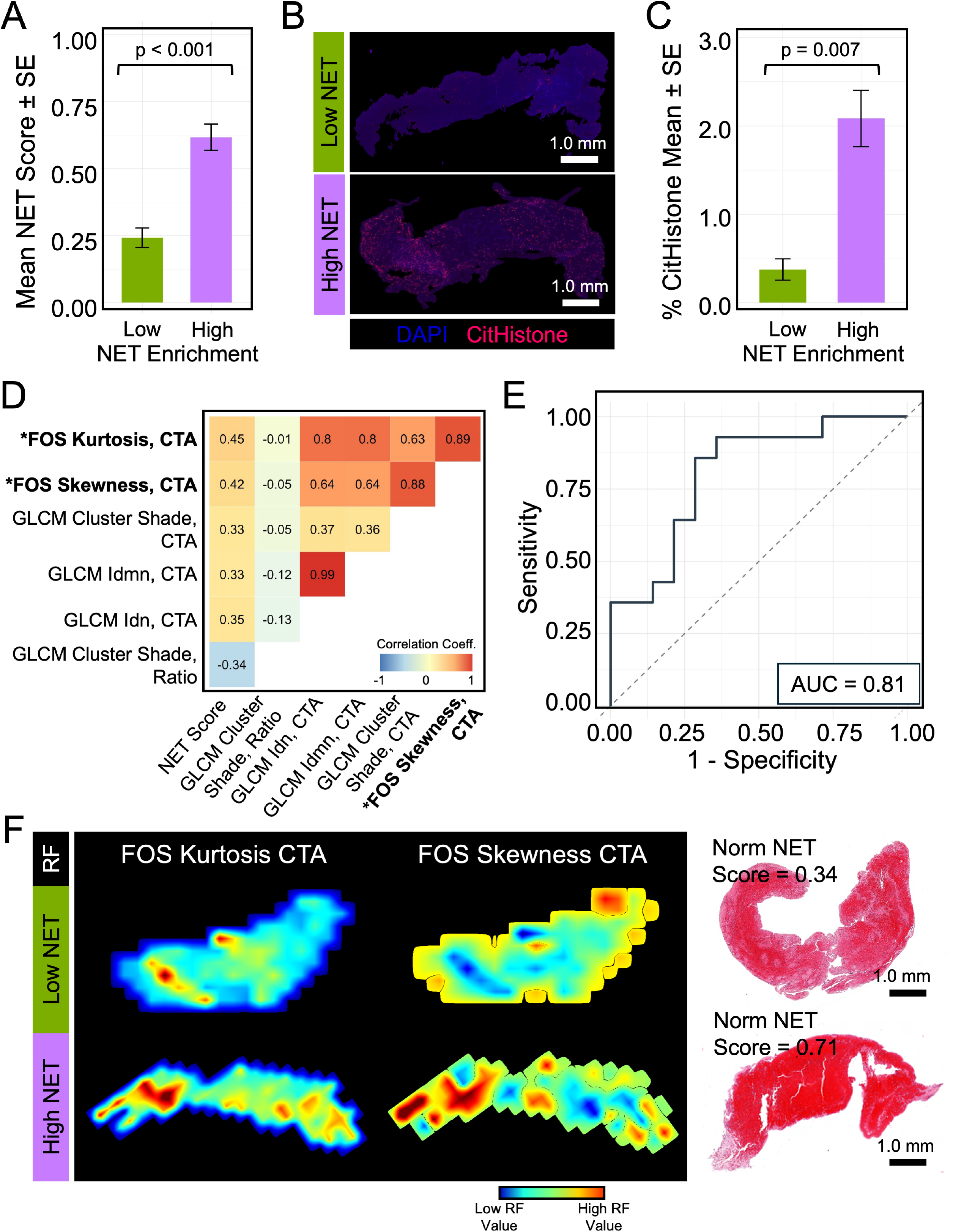
1: NET-enriched clots demonstrate a distinct radiomic signature associated with mechanical thrombectomy first pass outcome. **A)**. Bar plot shows significant difference between mean NET score for clots found to be low and high NET enriched. **B)**. Immunofluorescence images show whole-slide scans of clots stained for citrullinated histone, a marker of neutrophil extracellular trap formation, and DNA (DAPI). **C)**. Bar plot shows that clots classified as high NET enrichment based on transcriptomic analysis also demonstrated significantly increased percentage citrullinated histone by immunofluorescence quantification. **D)**. Heatmap shows correlation of clot radiomics with neutrophil extracellular trap formation score, Plotted are radiomics with a R>0.3 and p<0.1. Asterisks indicate P<0.05. **E)**. Receiver operating characteristic curve demonstrates an AUC of 0.81 for a multivariable logistic regression model predicting NET score based on the radiomics in D alone. **F)**. Radiomic heatmaps demonstrate differences in the CTA presentation of clots with low and high NET enrichment. Whole-slide digital pathology for the same clots is provided. Scalebars measure 1mm. (Abbreviations: NET=neutrophil extracellular trap; SE=standard error; CitHistone=citrullinated histone; %=percentage; FOS=first order statistic; GLCM=gray-level co-occurrence matrix; AUC=area under the receiver operating characteristic curve; RF=radiomic feature; Norm=normalized)

#### Identification of a radiomic signature for NET enrichment

Following validation of NET enrichment, we aimed to identify those radiomics correlated with NET scores. Overall, we identified 6 RFs demonstrating moderate correlation with NET score (p<0.10, R>0.3). Of these RFs, those with significant correlations to NET score included FOS Kurtosis (p=0.011) and FOS Skewness (p=0.019), both computed from CTA. A multivariable logistic regression model based on these 6 RFs was able to predict NET score with an AUC of 0.81 suggesting that pre-thrombectomy CT radiomics may be able to predict clot NET enrichment (Figure 3E). When two significant radiomic features (RFs) were visualized across both low and high NET groups, we observed elevated FOS Kurtosis and Skewness in clots with high NET enrichment (Figure 3F). High kurtosis reflects a predominance of similarly valued voxel intensities interspersed with an increasing number of extreme outliers, while high skewness indicates an asymmetric intensity distribution. In the context of acute ischemic stroke, a concurrent increase in both metrics suggests a clot composition characterized by dense tissue containing pockets of hyperintense regions. Corresponding histological analysis revealed that increasing NET enrichment was associated with a red blood cell–rich core, surrounded by a peripheral rim or shell of fibrin– platelet meshwork (Figure 3F). This histological pattern found in NET-rich clots aligns with the pro-thrombotic NET mechanism, promoting clot stability by facilitating RBC aggregation and retention, supporting fibrin cross-linking, and reducing clot permeability and breakdown.^15,34^

## Discussion

In this study we delved into the molecular underpinnings of AIS clots, with a focus on identifying biological processes associated with successful first-pass mechanical thrombectomy. Through transcriptomic profiling of extracted clots, we identified NET formation and the NET signaling pathway as the most significantly enriched processes linked to first-pass outcome. These findings highlight a critical role for NETs in influencing early thrombectomy outcomes and are consistent with prior studies.^6,30,32,34^

Our understanding of the role of NETs in AIS clot microstructure, thrombolytic resistance, and thrombectomy failure is evolving. Several prior *in vitro* studies have demonstrated that NET-rich clot analogs resist both thrombolysis and mechanical retrieval.^17,18^ Additional studies have shown that combination therapies such as tPA-DNase I, which target fibrin-platelet and DNA components, improve clot busting capabilities.^32,35^ Such findings have helped spur clinical trials investigating whether therapeutics with DNase I activity can improve reperfusion rates in large vessel occlusion stroke. In addition to these studies, several published works have demonstrated successful localization and quantification of NETs in retrieved thrombi. In a work by Ducroux et al.^32^, it was shown that NETs are constitutively present in AIS thrombi, with concentration in the outer layers. Moreover, it was found that thrombus NET content correlated significantly with the length of MT procedures and the number of device passes required. This study also expanded upon prior *in vitro* findings demonstrating that *ex vivo*, recombinant DNase I accelerated tPA-induced thrombolysis in NET rich clots.

Additional studies have investigated whether variation in clot NET enrichment is associated with stroke etiology (the origin of the clot) or clinical outcomes. In a work by Jabrah et al.^31^, it was found that cardioembolic clots had increased enrichment of neutrophils and extracellular web-like NETs compared to atherothrombotic or cryptogenic etiologies. A significantly higher distribution of web-like NETs was also observed around the clot periphery, a finding consistent with prior studies. In another recent work by Lapostolle et al.^30^, NET-rich thrombi were associated with unsuccessful recanalization and longer procedure time, as well as higher rates of post-operative neurological deficit (measured by changes in NIHSS) and disability or death (measured by modified Rankin Scale scores).

Despite growing recognition of the role of NETs in thrombectomy outcomes, no imaging-based metric currently exists to estimate clot NET enrichment from pre-procedural CT. Developing such a metric is critical for personalized medicine, e.g., stratifying patients by their likelihood of benefiting from combination therapy with tPA and DNase I. Non-invasive identification of NET-rich clots could inform personalized treatment strategies, improve reperfusion rates, and enhance overall thrombectomy success. Therefore, given our transcriptomic results, we chose to further investigate whether clot RFs extracted from pre-thrombectomy CT could be used to potentiate a non-invasive digital biomarker of NET enrichment and thus first-pass success.

We first set out to identify radiomics significantly different between mFP outcomes and found 40 significant RFs. These RFs, computed from CTA and nCCT included several RFs previously reported in the context of AIS.^36–38^ Additionally, several of these RFs, such as FOS Energy, were identified in prior work as scale-invariant RFs correlating well with WBC percent composition.^33^ Given these findings, we next investigated if this same set of RFs also demonstrated correlation with clot NET enrichment. To achieve this, we enlisted existing bioinformatics techniques and databases to derive an estimate of single-sample gene set enrichment for NET formation. Recognizing that our computed NET enrichment metric was inferred from clot transcriptomes, we further validated our NET score using ground truth immunofluorescence labeling and whole-slide quantification of citrullinated histone, an established marker of NET formation.^30–32,39,40^

From comparison of RFs between mFP outcomes we found that 40 RFs demonstrated significant difference. Upon further correlation with clot NET score, we identified a 6 RF signature associated with NET enrichment which in multivariable logistic regression achieved an AUC of 0.81. Using immunofluorescence staining and whole-slide digital pathology quantification we further validated our NET score metric. This metric allowed us to establish a direct link between the thrombus NET enrichment and specific RFs, helping to unravel the intricate interplay between molecular and imaging characteristics in the context of MT first pass success.

Our study provides a nuanced understanding of the complex biological factors influencing mechanical thrombectomy success. Several limitations should be noted. First, the sample size was relatively small, which may limit the generalizability of our findings. Validation in larger, multi-center cohorts with diverse patient populations is needed to confirm the robustness and clinical utility of the identified transcriptomic and radiomic signatures. Second, although our radiomic analysis demonstrates potential for non-invasive prediction of NET enrichment, these findings are based on retrospective imaging data and require prospective validation. Lastly, the influence of stroke etiology, clot age, and pre-hospital treatments on NET-related signatures warrants further investigation. Nonetheless, our study is among the first to integrate clot transcriptomics with CT radiomics and digital pathology to identify a specific prothrombotic mechanism, NET enrichment, implicated in MT outcomes.

## Conclusions

Our findings highlight the critical role of NET enrichment in influencing first-pass mechanical thrombectomy outcomes in acute ischemic stroke. By integrating clot transcriptomics with radiomic analysis of pre-thrombectomy CT imaging, this is the first study to identify a non-invasive signature of NET-rich thrombi for stroke. This approach lays the groundwork for stratifying patients prior to intervention, enabling tailored therapeutic strategies such as the early use of combination tPA-DNase I therapy or optimized device selection. With further validation in larger, prospective cohorts, radiomic assessment of NET content may become a valuable clinical tool to improve reperfusion rates and overall stroke outcomes.

## Acknowledgements

We acknowledge the assistance of the Multispectral Imaging Suite and Histology Core Laboratory in the Department of Pathology & Anatomical Sciences, Jacobs School of Medicine and Biomedical Sciences, University at Buffalo.

## Disclosures

BAS – None

TRP – None

SMMJS – None

KEP – Ownership: Neurovascular Diagnostics, Inc.

SB – None

AS – None

TDJ – None

EIL – Board Membership: Stryker, NeXtGen Biologics, MedX Health, Cognition Medical, EndoStream; Consultancy: Claret Medical, GLG Consulting, Guidepoint, Imperative Care, Medtronic, Rebound Therapeutics, StimMed; Employment: University at Buffalo Neurosurgery Inc; Expert Testimony: renders medical/legal opinions as an expert witness; Stock/Stock Options: NeXtGen Biologics, Cognition Medical, Rapid Medical, Claret Medical, Imperative Care, Rebound Therapeutics, StimMed.

AHS – Financial Interest/Investor/Stock Options/Ownership: Adona Medical, Inc., Amnis Therapeutics, Bend IT Technologies, Ltd., BlinkTBI, Inc, Buffalo Technology Partners, Inc., Cardinal Consultants, LLC, Cerebrotech Medical Systems, Inc, Cerevatech Medical, Inc., Cognition Medical, CVAID Ltd., Endostream Medical, Ltd, Imperative Care, Inc., Instylla, Inc., International Medical Distribution Partners, Launch NY, Inc., NeuroRadial Technologies, Inc., Neurotechnology Investors, Neurovascular Diagnostics, Inc., PerFlow Medical, Ltd., Q’Apel Medical, Inc., QAS.ai, Inc., Radical Catheter Technologies, Inc., Rebound Therapeutics Corp. (Purchased 2019 by Integra Lifesciences, Corp), Rist Neurovascular, Inc. (Purchased 2020 by Medtronic), Sense Diagnostics, Inc., Serenity Medical, Inc., Silk Road Medical, SongBird Therapy, Spinnaker Medical, Inc., StimMed, LLC, Synchron, Inc., Three Rivers Medical, Inc., Truvic Medical, Inc., Tulavi Therapeutics, Inc., Vastrax, LLC, VICIS, Inc., Viseon, Inc. Consultant/Advisory Board: ational Center for Advancing Translational Sciences of the National Institutes of Health under award number UL1TR001412 to the University at Buffalo.

JK – None.

VMT – Financial Interest/Investor/Stock Options/Ownership: Neurovascular Diagnostics, Inc., QAS.ai, Inc. Grant Support: Brain Aneurysm Foundation, National Science Foundation, NIH NINDS, clinical and translational science institute grant from the national Center for Advancing Translational Sciences of the National Institutes of Health under award number UL1TR001412 to the University at Buffalo. Consultant/Advisory Board: Canon Medical Systems America.

